# Using the AKAR3-EV biosensor to assess Sch9- & PKA-signalling in budding yeast

**DOI:** 10.1101/2022.10.27.514151

**Authors:** Dennis Botman, Sineka Kanagasabapathi, Bas Teusink

**Author notes:** These authors contributed equally to this work (Dennis Botman, Sineka Kanagasabapathi). Correspondence should be addressed to Bas Teusink.

## Abstract

Budding yeast uses the well-conserved TORC1-Sch9 and cAMP-PKA signalling pathways to regulate adaptations to changing nutrient environments. Dynamic and single-cell measurements of the activity of these two cascades will improve our understanding of cellular adaptation of yeast. Here, we employed the AKAR3-EV biosensor developed for mammalian cells to measure the cellular phosphorylation status determined by Sch9 and PKA activity in budding yeast. Using various mutant strains and inhibitors, we show that AKAR3-EV robustly measures the Sch9- and PKA-dependent phosphorylation status in intact yeast cells. At the single-cell level, we found that the phosphorylation responses are homogenous for glucose, sucrose and fructose, but highly heterogeneous for mannose. The Sch9 and PKA pathways have a relatively high affinity for glucose (K_0.5_ of 0.24 mM) under glucose derepressed conditions. Lastly, steady-state FRET levels of AKAR3-EV seem to be independent of growth rates, suggesting that the Sch9- and PKA-dependent phosphorylation activity are transient responses to nutrient transitions. We believe that the AKAR3-EV sensor is an excellent addition to the biosensor arsenal for illuminating cellular adaptation in single yeast cells.

## Introduction

One universal aspect of life is change, and the ability to adapt to it is a major determinant of reproductive success. For the unicellular organism *Saccharomyces cerevisiae* (budding or baker’s yeast), there is no exception. In the wild, this yeast lives on fruit and tree barks where it endures feast-famine cycles^1,2^. In industry, domesticated yeast also experiences changing conditions when used in large-scale fermenters with inoculation transitions and poor stirring^3,4^. For budding yeast, nutrient availability is a major environmental parameter that sets the investment in metabolism, stress resistance and proliferation^5,6^. Understanding the logic of these circuitries is a key challenge in cell biology. Also for industry, control over this response, e.g. by selection of certain preferred subpopulation or removal of undesired populations, could increase production efficiencies^7^. Nutrient adaptations can be captured on population level using bulk assays, but these methods may mask the true responses produced by the intracellular signalling circuits inside single cells. Thus, a more profound characterisation of cellular adaptation of budding yeast to nutrient changes at the single-cell level is highly desired.

Regarding carbon sources, fruit sugars are preferred by yeast and therefore it has developed various pathway to sense and adapt to changes in the availability of these substrates^8–10^. The cAMP-PKA pathway is one of the major signalling cascades which get activated when cells encounter fermentable sugars ^11–13^. cAMP production is activated via two routes^11,14–29^: via import and metabolism of the sugar (1) and via extracellular sensing of glucose and sucrose by the G-protein coupled receptor Gpr1 (2). These 2 inputs give a transient increase in cAMP, which causes dissociation of Bcy1 (a PKA regulator) from the PKA subunits Tpk1-3. This finally results in an increase of PKA activity; PKA is a major effector kinase in yeast, accounting for 75-90% of the cellular changes during a transition from a glucose derepressed (respiratory) state to a fermentable glucose repressed (fermentative) state^5,8,9,30–33^. The evoked transition gives a radical change in yeast physiology; cells change their metabolism from respiratory to fully fermentative, repress metabolic pathways for other carbon sources, decrease their stress resistance and make large investments in ribosomal biogenesis.

The TORC1-Sch9 cascade exerts a second major signalling cascade in yeast cells^34–37^. In contrast to PKA, which get activated by mostly fermentable sugars, Sch9 is activated by TORC1 when a complete palette of nutrients for growth (such as amino acids, nitrogen, phosphate and a fermentable carbon source) are available^34,36,38.31,36^. Although the two pathways can operate independently^39^, they have many positive interactions and can rescue each others activities, probably via the large overlap in their targets: Sch9 and PKA both phosphorylate the RRXT motif ^40–43^. Activation of Sch9 activates 90% of the genes that PKA also activates^32^, making Sch9 as important as PKA for proper cellular decision making in yeast.

Currently, PKA and Sch9 activities are difficult to measure in (single) cells: the most common method is measuring the activity of trehalase, a PKA and Sch9 target, or using kemptide as a substrate^35,36^. However, these bulk assays lack single-cell information and show only static activity levels of PKA and Sch9 activity. Studies suggest that the PKA activity in yeast is more a transient phenomenon activated during (mostly) sugar transitions and that TORC1-Sch9 axis dictates the steady-state growth mode of a yeast cell^36,44^. Dynamic readouts are needed to substantiate these interesting hypotheses. Moreover, single-cell dynamics allows to test for heterogeneity during nutrient transitions – a phenomenon that we did not find for cAMP dynamics,^13^ but it is unknown if heterogeneity exists more downstream. Furthermore, the sensitivity of Sch9 and PKA activity with respect to glucose remains to be characterised. Finally, the basal RRXT phosphorylation status of the cell at various growth rates is also poorly characterized.

Here, we implemented and tested the mammalian PKA sensor AKAR3-EV in yeast, to provide a tool that can help to enlarge our understanding of PKA and Sch9 signalling. The sensor allowed us to measure the single-cell dynamics of PKA and Sch9 phosphorylation status in a robust and accurate manner. We found large heterogeneous responses of yeast cells for some nutrient transitions which we did not previously find in cAMP dynamics. The detected heterogeneity potentially affects the overall cellular state since these two kinases constitute the vast majority of the cellular transition during a transition to a fermentable carbon source. Furthermore, our data implies that the phosphorylation status is not related to growth rate. How the two kinases regulate the cellular transitions at a single-cell level can be studied in more depth using this sensor.

## Material & Methods

### Sensor construction

AKAR3EV with YPET-eCFP as FRET pair was kindly provided by dr. Aoki^45^. The sensor was amplified using KOD One™ PCR Master Mix (Toyobo, Osaka, Japan) with 5′-ATGCTAGCACGGAGCTCACTGAATTCGGCATGGTGAG-3′ and 5′-ATGGATCCACGGTCGACACTTTAATCCAGAGTCAGGCG-3′ as forward and reverse primer, respectively. Next, the PCR product and the pDRF1-GW plasmid were digested using BamHI-HF and NheI-HF (New England Biolabs, Ipswich, MA) and the PCR product was ligated into the plasmid using T4 DNA ligase (New England Biolabs), which yield pDRF1-GW AKAR3-EV.

AKAR3-EV-NR was constructed by performing a PCR with KOD One™ PCR Master Mix on pDRF1-GW AKAR3-EV with 5′-TATTCCGGATTGAGGCGCGCGGCCCTGGTTGACGGCGGCCGCATGGTGAGCAAGGGC-3′ as forward primer and 5′-ATGGATCCACGGTCGACACTTTAATCCAGAGTCAGGCG-3’ as reverse primer. The PCR product and pDRF1-GW AKAR3-EV were digested using Kpn2I and XbaI (Thermo Fisher Scientific, Waltham, MA, USA). Afterwards, the PCR product was ligated in the digested pDRF1-GW AKAR3-EV, replacing the sensor domain and eCFP with the non-responding (T506A) sensor domain and eCFP. pDRF1-GW eCFP was made by performing a PCR with KOD One™ PCR Master Mix on AKAR3-EV with FW primer 5′-ATGCTAGCATGGTGAGCAAGGGCG-3′ and RV primer 5′-TAGCGGCCGCTTACTTGTACAGCTCGTCCATGCCG −3′ after which the PCR product and pDRF1-GW were digested using NheI-HF and NotI-HF (New England Biolabs). Finally, the PCR product was ligated into pDRF1-GW using T4 DNA ligase.

### Yeast transformation

Strains used in this study are listed in table 1. Strains were transformed by resuspending yeast cells from either a YPAD plate or a selective plate in a transformation mixture containing 240 μL PEG 3350 (50%w/v), 40 μL 1M LiAC, 10 μL Salmon Sperm DNA (10 mg/mL, Sigma Aldrich, Stl. Louis, MO, USA) and 500-1000 ng plasmid DNA. Next, cells were incubated for 15-20 minutes at 42°C. Afterwards, the cells were centrifuged at 13,000 g for 30 seconds, supernatant was removed and 150 μL water was added. Finally, the cells were plated on selective plates.

**Table 1.**
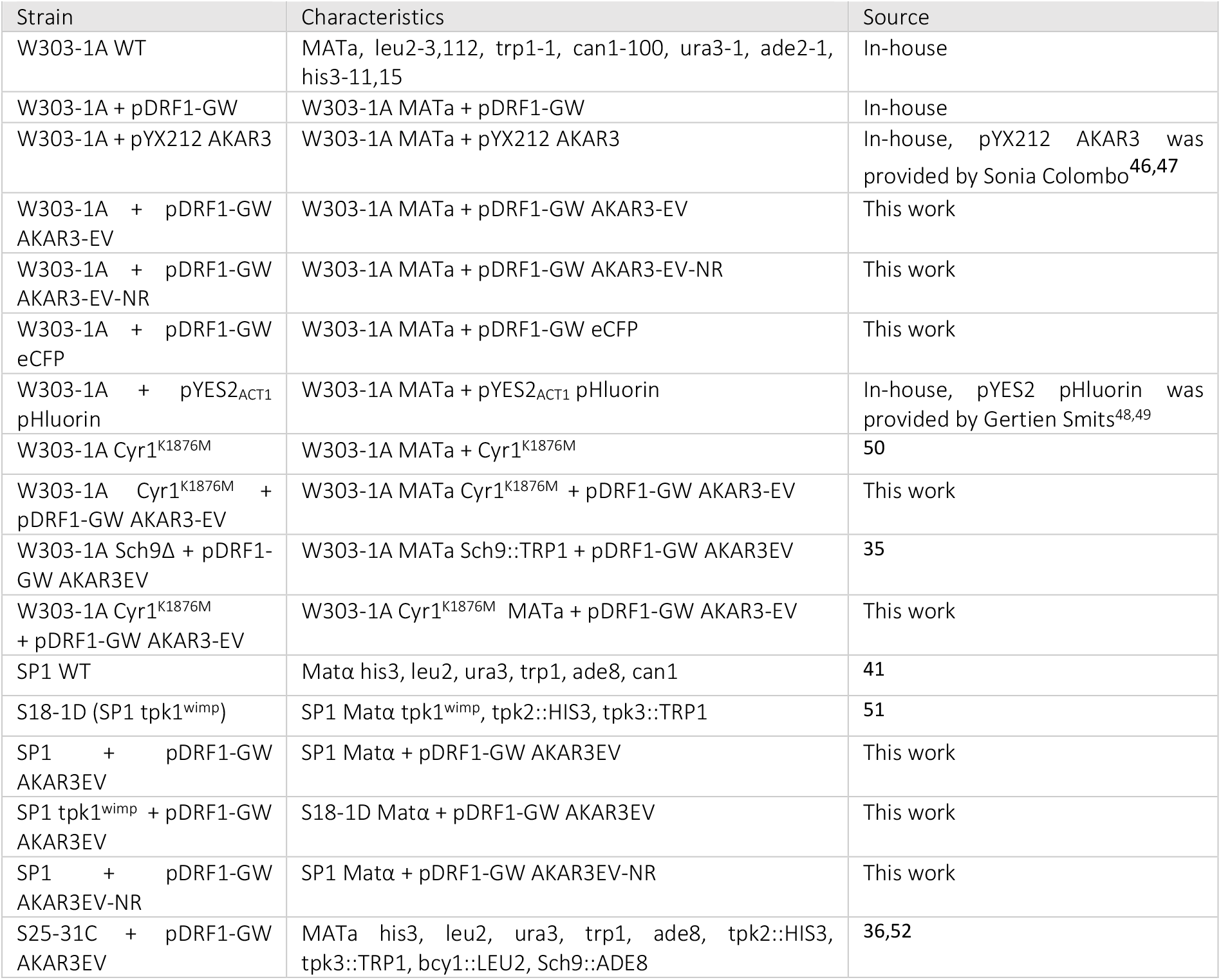
Used strains in this study.

### Yeast growth

Cells expressing URA3 plasmids were grown overnight in 1× YNB medium (Sigma Aldrich, St. Louis, MO, USA), containing 1% Ethanol (Boom BV, Meppel, Netherlands), 20mg/L adenine hemisulfate (Sigma-Aldrich), 20mg/L L-tryptophan (Sigma-Aldrich), 20mg/L L-histidine (Sigma Aldrich) and 60mg/L L-leucine (SERVA Electrophoresis GmbH, Heidelberg, Germany). For WT strains, uracil (Sigma Aldrich) was added to a final concentration of 20mg/L. The cells were subsequently diluted and grown overnight to an OD_600_ of 0.1-1.5, with at least 5 divisions.

For the experiments that involved S25-31C, cells were grown on 1× YNB medium containing 1% Ethanol, 5 mM glucose (Boom BV, Meppel, The Netherlands), 20mg/L adenine hemisulfate, 20mg/L L-tryptophan, 20mg/L L-histidine and 60mg/L L-leucine until glucose was exhausted. Next, cells were kept on this medium for 2 more days after which they were visualized under the microscope.

### ConA coverslips

Concanavalin (ConA) coverslips were made as described by Hansen et al., 2015^53^. To prepare the coverslips, the ConA was diluted to 200 μg/mL and put on coverslips. The coverslips were dried overnight in a 6 wells plate.

### Microscopy

Cells were grown as described and transferred to the 6 wells plates containing the ConA-coated coverslips. Next, the coverslip was placed in a Attofluor cell chamber (ThermoFisher Scientific, Waltham, MA) and 1 mL of medium was added to the cell chamber. The coverslips were visualized using a Nikon Ti-eclipse microscope (Nikon, Minato, Tokio, Japan) at 30°C equipped with a TuCam system (Andor, Belfast, Northern Ireland) and 2 Andor Zyla 5.5 sCMOS Cameras (Andor) and a SOLA 6-LCR-SB light source (Lumencor, Beaverton, OR). Cells expressing the sensors, except pHluorin, were excited via a 438/24 nm excitation filter (Semrock, Lake Forest, IL) and emission was passed through a 458 nm long-pass (LP) dichroic mirror. The acceptor and donor emissions were filtered by a 542/27 nm and 483/32 nm filter (Semrock). Direct acceptor fluorescence was recorded with a 500/24 nm excitation filter, a 520 LP dichroic filter and a 542/27 nm emission filter. pHluorin was excited at 460-500 nm and 380-420 nm and emission was recorder at 510-560 nm. For all perturbations, a baseline was recorder after which YNB medium containing the compound of choice in a 10x concentration was added and diluted in the cell chamber to a 1x concentration.

### Rapamycin experiments

Cells were grown as described and incubated with 200 nM rapamycin or a solvent (100% Ethanol) for at least 2 hours after which the perturbations (addition of 10 mM glucose) were performed.

### Microscopy Analysis

Cells were segmented using an in-house script. In brief, this script stabilizes any drift using the image stabilizer plugin^54^. Next, background correction was performed and cells were segmented using the Weka Segmentation plugin^55,56^ and the mean fluorescence for each cell was calculated for each frame. The resultant text files were analysed using R 4.1.3. For all cells, 40% bleedtrough correction was performed and the FRET ratio (i.e. bleedtrough-corrected YFP divided by CFP signal) was calculated. Finally, baseline normalization was performed for time-lapse data. pHluorin ratios were calculated by dividing the fluorescence at 380-420 nm excitation over the fluorescence at 460-500 nm.

For the dose-response fit, final FRET levels (i.e. the mean FRET value of the last 3 frames) after the glucose additions were fitted (using the nls function in R) against the final glucose concentration according equation 1, with [glucose] as the glucose concentration in mM, max as the maximal change in normalized FRET and K_0.5_ as the glucose concentration giving 50% of the maximal response.

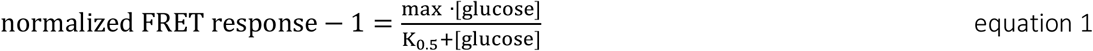

Clustering was performed using the factoextra package in R. The optimal amount of clusters was determined by eye.

### Flow cytometry

Cells were grown as described with YNB medium containing either 1% Ethanol, 100 mM glucose, 100 mM galactose or 100 mM mannose. Next, samples were measured using a CytoFLEX S Flow Cytometer (Beckman Coulter, Brea, CA, United States). Cells were excited using a 405 nm and a 488 nm laser and emission fluorescence was passed through a 470/20 and 525/40 nm filter and recorded by avalanche photodiodes. Events with a saturating forward or side scatter were filtered after which the median fluorescence signal of the cells expressing empty pDRF1-GW plasmid was subtracted from all samples. Next, bleedtrough was calculated using the eCFP-expressing strain, and cells with at least a acceptor fluorescence signal of 2500 (arbitrary units) were kept. Lastly, FRET ratios were calculated for all remaining cells.

### Growth assays

Yeast strains were grown on 1% Ethanol medium as described. Cells were washed twice by centrifuging at 3500g for 3 minutes and resuspending in YNB medium without carbon source. Afterwards, cells were resuspended in YNB medium without carbon source to an OD_600_ of 1. Next, 20 μL of cells were put in well of a 48-well microtiter plate having 480 μL YNB medium containing either 10 mM glucose, 10 mM galactose or 0.1% ethanol. OD_600_ was measured every 5 minutes with a CLARIOstar plate reader (BMG LABTECH, Ortenberg, Germany) at 30°C and 700 rpm orbital shaking. Growth rates were calculated by calculating a moving average over each growth curve. Next, a sliding window was used in which a linear regression was fitted for each window which gives the growth rate during this window. Next, the tenth fastest found slope was selected as the determined growth rate.

## Results

### AKAR3-EV shows a FRET response in budding yeast

To visualize cellular kinase activities of Sch9 and PKA, we tested the use of the AKAR3-EV sensor, which consists of YPET-FHA1-EVlinker-RRAT_motif_-eCFP. This sensor, developed for mammalian PKA assays, should also work in yeast as PKA and Sch9 also have RRXT as recognition site^40–42^. We constructed a non-responsive AKAR3-EV-NR sensor as a control by mutating the threonine in the RRXT motif to alanine (T506A), and expressed both AKAR3-EV and AKAR3-EV-NR in the W303-1A and SP1 strains. Both W303-1A and SP1 were used as some mutant W303-1A strains have difficulties to express sensors. This was also found for the yEPAC cAMP sensor and indicates a more general problem for some W303-1A mutants and not a sensor-specific issue.

In these two strains, we assessed the FRET response from a 1% ethanol to a 100 mM or a 10 mM glucose transition (Figs. 1A and 1B). Furthermore, we tested the older generation AKAR3 sensor for performance comparison^46,47^. The AKAR3-EV sensor gave a clear increase in FRET after glucose addition, which was 3-4 times higher compared to AKAR3-EV-NR. Furthermore, we found only a marginal response of the original AKAR3 sensor, showing that the AKAR3-EV sensor has an improved ability to visualize phosphorylation activity of PKA and Sch9 in yeast. Cellular expression was more than sufficient, and the sensor shows a uniform distribution in cells (Fig. 1C). Of notice, the AKAR3-EV-NR sensor shows a significant increase shortly after glucose addition in the W303-1A strain. This increase is not caused by osmotic changes since 100 mM sorbitol addition did not increase FRET levels (Fig. S1A). Conclusions about FRET responses of AKAR3-EV directly after a transition should therefore be taken carefully. Furthermore, the AKAR3-EV shows now basal drift in FRET which we found for AKAR3. The original AKAR3 sensor also shows a small dip in FRET response after the glucose addition, which could come from differential pH sensitivity of the fluorescent proteins in this sensor. AKAR3-EV uses YPET-eCFP as a FRET pair, which shows more pH robustness compared to the eCFP-Venus FRET pair used in AKAR3 (Fig. S1B)^57^. Lastly, expression levels of AKAR3-EV did not affect basal FRET levels and growth was not affected on various carbon sources (Figs. S1C and S1D). In conclusion, the AKAR3-EV can be used in yeast to assess cellular phosphorylation of the RRXT motif.

**Figure 1.**
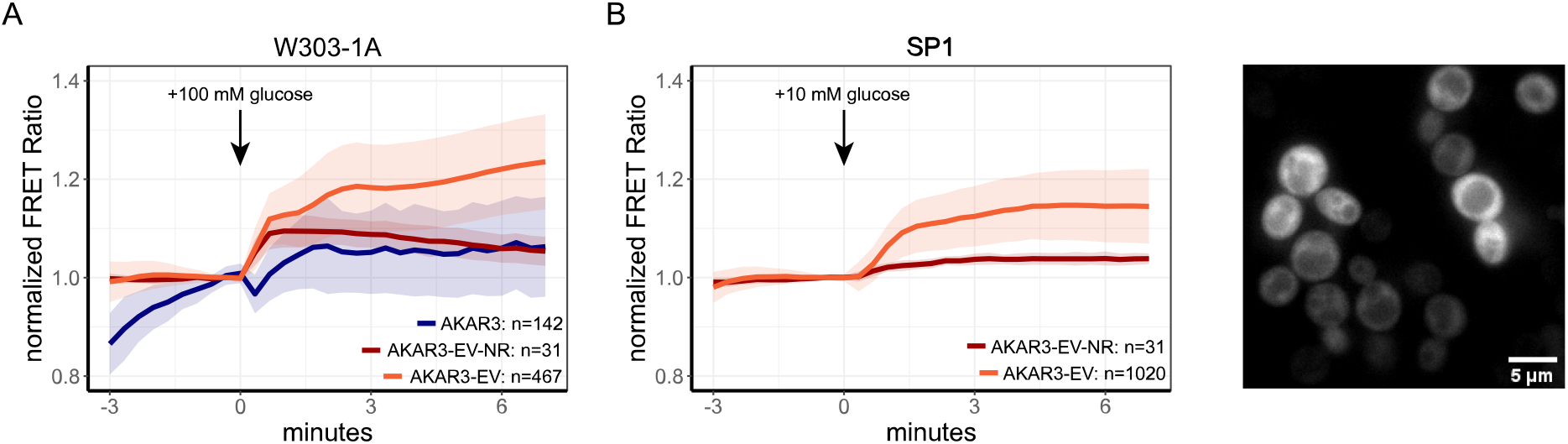
AKAR3-EV responds to glucose addition. A) Response of W303-1A WT cells expressing AKAR3, AKAR3-EV NR or AKAR3-EV grown on 1% ethanol to 100 mM glucose addition. B) Response of SP1 WT cells expressing either AKAR3-EV or AKAR3-EV NR grown on 1% ethanol to 10 mM glucose addition. C) Expression and distribution of W303-1A cells expressing AKAR3EV. Lines show mean FRET value, normalized to the baseline, shades indicate SD, colors indicate the expressed sensor. All data is obtained from at least 2 biological replicates.

### AKAR3-EV responses are dependent on Sch9 and PKA signalling

To assess the influence of cAMP-PKA and TORC1-Sch9 signalling on the FRET readout, we compared FRET responses of W303-1A and SP1 WT to strains carrying the tpk1^wimp^ mutation (which has low PKA activity) and the Sch9Δ strain upon a shift from 1% Ethanol to 10 mM glucose (Figs.2A & 2B). The tpk1^wimp^ shows a lower initial response but does gradually shifts to WT FRET levels at the end of the timelapse (normalized FRET levels of 1.16 and 1.14 for WT and tpk1^wimp^, respectively). In contrast, Sch9 deletion gave a reduction in the reached plateau of 25% (normalized FRET levels of 1.24 and 1.18 for WT and Sch9Δ, respectively). We also tested whether W303-1A Cyr1^K1876M^, which lacks the classical cAMP peak upon glucose addition shows an altered AKAR3-EV response. This strain showed a response that reached a plateau similar to the sch9Δ strain (Fig. 2C), but with different dynamics.

**Figure 2.**
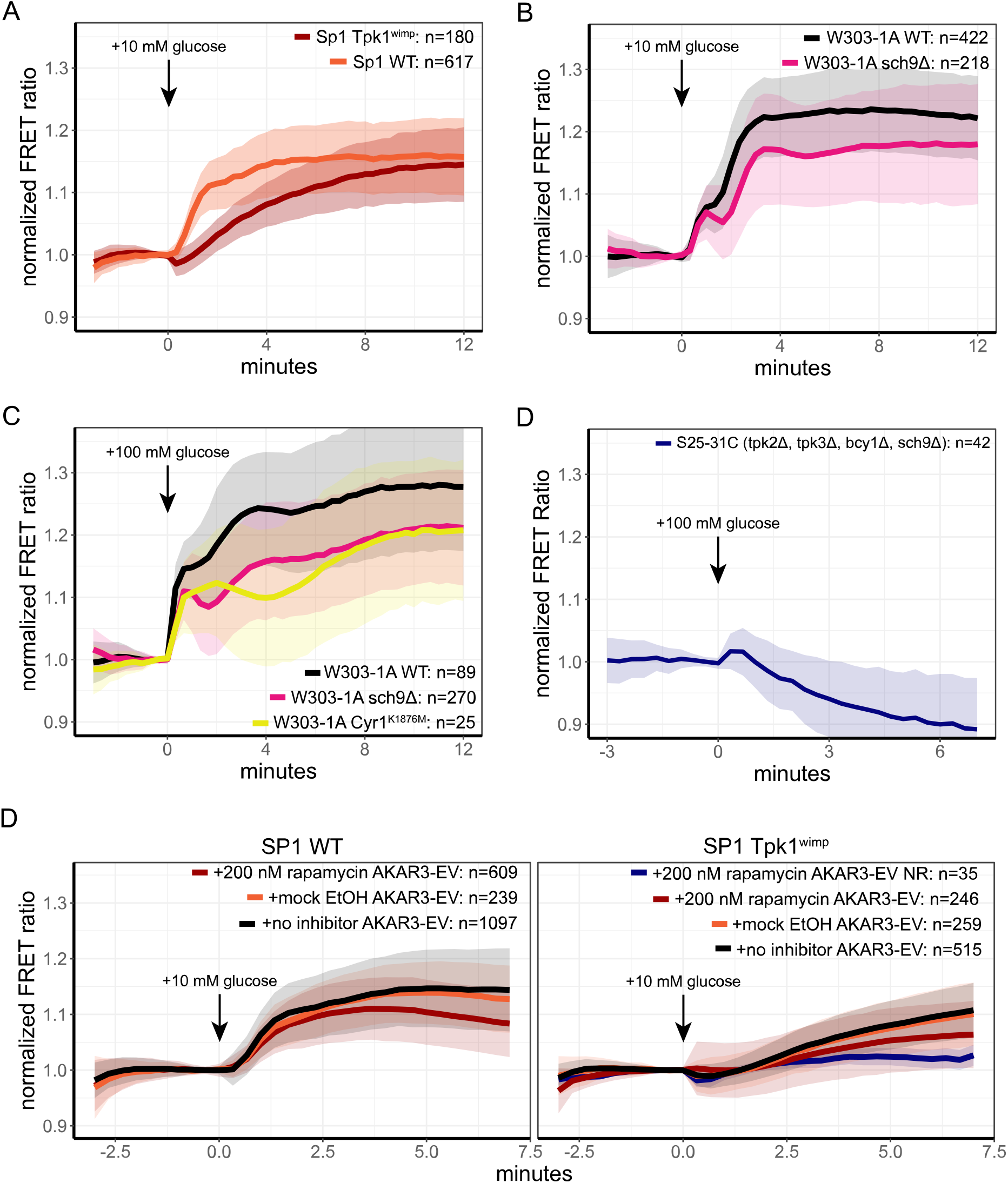
AKAR3EV responds to both Sch9 and PKA activity. A) FRET response of SP1 WT and SP1 Tpk1^wimp^ cells expressing AKAR3EV grown on 1% Ethanol to 10 mM glucose addition. B) FRET response W303-1A WT and W303-1A sch9Δ cells expressing AKAR3EV grown on 1% Ethanol to 10 mM glucose addition. C) FRET response W303-1A WT, W303-1A sch9Δ and W303-1A Cyr1^K1876M^ cells expressing AKAR3EV grown on 1% Ethanol to 100 mM glucose addition. D) FRET response of S25-31C cells expressing AKAR3EV grown on 1% Ethanol to a 100 mM glucose pulse. D) FRET response of SP1 WT and SP1 Tpk1^wimp^ cells expressing AKAR3EV or AKAR3-EV NR to 10 mM glucose addition. Cells were grown on 1% Ethanol, incubated for at least 2 hours with rapamycin, mock (ethanol) or without any addition and pulsed with 10 mM glucose. Lines show mean FRET value, normalized to the baseline, shades indicate SD, colors indicate the expressed sensor, strain, and/or treatment. All data is obtained from at least 2 biological replicates.

Since both cAMP-PKA and TORC1-Sch9 cascades affect the AKAR3-EV response, we assessed whether a strain mutated in both cascades showed any increase in FRET levels upon glucose addition. For this we used the S25-31C strain (tpk2Δ, tpk3Δ, bcy1Δ, Sch9Δ) which is known to have impaired phosphorylation activity assessed by trehalase activity^36^. As expected, 100mM glucose addition to this strain showed no increase in FRET levels of AKAR3-EV, indicative that AKAR3-EV FRET (and hence RRXT phosphorylation) levels rely on PKA and Sch9 signalling (Fig. 2D). In contrast, there is a slight decrease in FRET signal, which may be explained by glucose-induced phosphatase activity on the RRXT motif – the phosphorylation state of the sensor is in the end a steady-state balance between phosphorylation and dephosphorylation activity.

Finally, we confirmed the dependency of RRXT phosphorylation on Sch9 and PKA by simultaneous suppression of both pathways. For this, we used the TORC1 inhibitor rapamycin to inhibit this signalling cascade and combine this with the SP1 WT and tpk1^wimp^ strains which has decreased (but not completely absent) PKA activity. In wild-type SP1, we found a decrease of 27% in FRET response of rapamycin-treated cells compared to untreated cells after a 10 mM glucose pulse (Fig. 2D). SP1 tpk1^wimp^ cells already showed a 27% decreased maximal response (compared to WT cells) which further decreased to 60% when treated with rapamycin. This response was still higher compared to the response of the non-responsive sensor, which most likely is attributed to the remaining activity of tpk1^wimp^.

In conclusion, we show that the AKAR3-EV sensor can be used to measure TORC1-Sch9 and cAMP-PKA activity by visualizing the cellular RRXT phosphorylation status. Deletion of Sch9 or removal of the classical cAMP peak results in a decreased phosphorylation status of the cell. In contrast, decreased PKA activity gives a decreased initial response, but eventually reaches the same basal phosphorylation levels as wild-type cells.

### Sugar transitions shows distinct phosphorylation dynamics with single-cell heterogeneity

In our previous study we found that cAMP signalling in yeast is different for different sugars, and it was homogeneous with no clear subpopulations or non-responders^13^. We used the AKAR3-EV sensor to measure the downstream response to different sugars. W303-1A cells grown on 1% Ethanol medium were pulsed with either 100 mM glucose or sucrose (both Gpr1 agonists), 100 mM fructose (no Gpr1 agonist or antagonist) or 100 mM mannose (an antagonist of Gpr1)^13,58^. Glucose, sucrose and fructose gave a clear transient increase in FRET ratio after addition, where sucrose and glucose gave a slightly faster and more sustained response. In contrast, mannose showed a decline in FRET signal after which the FRET response increased to higher levels which sustained until the end of the timelapse recording (Fig. 3A). For fructose, glucose and sucrose, we found no clear and significant subpopulations. The most striking response at single-cell level, however, was found for mannose addition. Mannose addition gave a highly heterogenous response (Figs. 3B-3E, movie S1) which was not found for the non-responsive sensor (Fig. S2). After the dip in FRET levels (which all cells do seem to have), we identified 3 discrete responses, which we clustered using k-means clustering. The first cluster consisted of cells that have a broad timeframe in which cells increase in FRET levels. Furthermore, final FRET levels are slightly increased (mean normalized FRET value of 1.08). The second cluster showed clear switchers which obtained a final normalized FRET value of 1.25. Finally, the third cluster consisted of cells that did not increase but decreased their RRXT phosphorylation levels (mean normalized FRET value of 0.90 at the end of the timeframe). Interestingly, for the mannose transitions, cells that had a lower baseline FRET value also seem to end in a lower FRET state and vice versa (Fig. 3D). In conclusion, the AKAR3-EV sensor shows multifarious responses after sugar addition, dependent on the specific sugar. For mannose, we found a highly heterogenous response with 3 subpopulations: committers, doubters and stayers.

**Figure 3.**
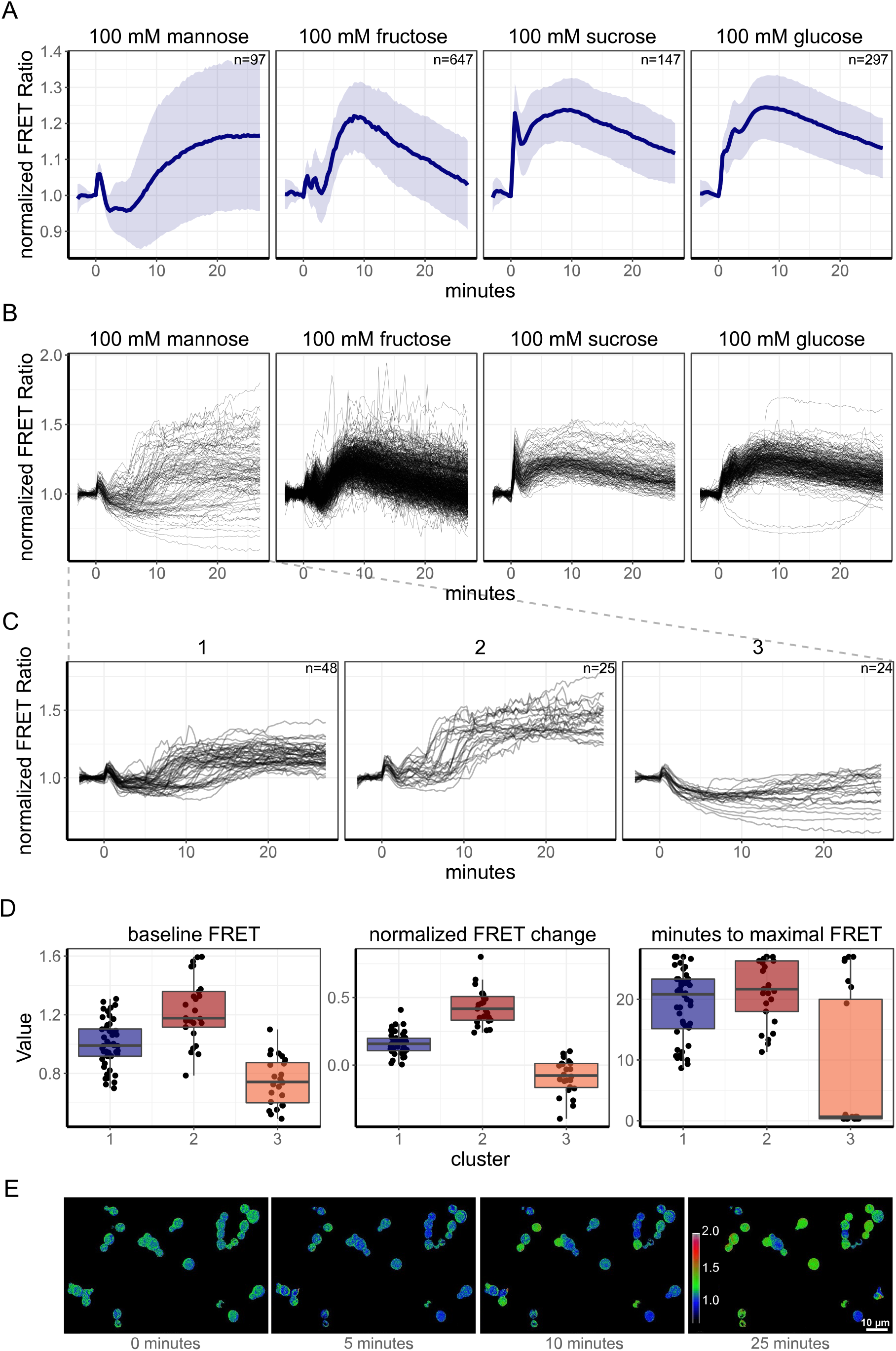
Single-cell assessment of AKAR3EV responses shows heterogeneity for mannose, but not for fructose, sucrose and glucose additions. A) Response of W303-1A WT cells expressing AKAR3EV grown on 1% ethanol to the various sugar additions, added at t=0 minutes. Lines show mean FRET value, normalized to the baseline, shades indicate SD. Facet titles show the final concentration of the added sugar. B) Baseline-normalized single-cell traces of the transitions depicted in A. C) The 3 identified clusters found for the heterogenic response after 100 mM mannose addition. Lines depict single-cell traces of the normalized FRET value. D) Basal FRET level (not baseline normalized), the change in normalized FRET values at the end of the recorded timelapse and the timing of the maximal obtained FRET value for each cluster. Each dot depicts a single cell. Boxplots depict median with quartiles; whiskers indicate largest and smallest observations at 1.5 times the interquartile range. All data is obtained from at least 2 biological replicates. E) Static pictures of a 1% Ethanol to 100 mM mannose transition. Colors indicate FRET ratio (normalized to the first baseline frame), mannose was added at 0 minutes. Scale bar indicates 10 μm.

### The nutrient-induced phosphorylation system has a high affinity for glucose

When we tested the response of the ACAR3-EV sensor in W303-1A to different levels of glucose (Fig 4A), we found a graded response that could be fitted by a single binding curve with a relatively high affinity, with a K_0.5_ of 0.27 mM for glucose (Fig. 4C). In addition, no clear subpopulations or non-responders were detected (Fig. S3). This is in line with previous results found for cAMP^13^.

**Figure 4.**
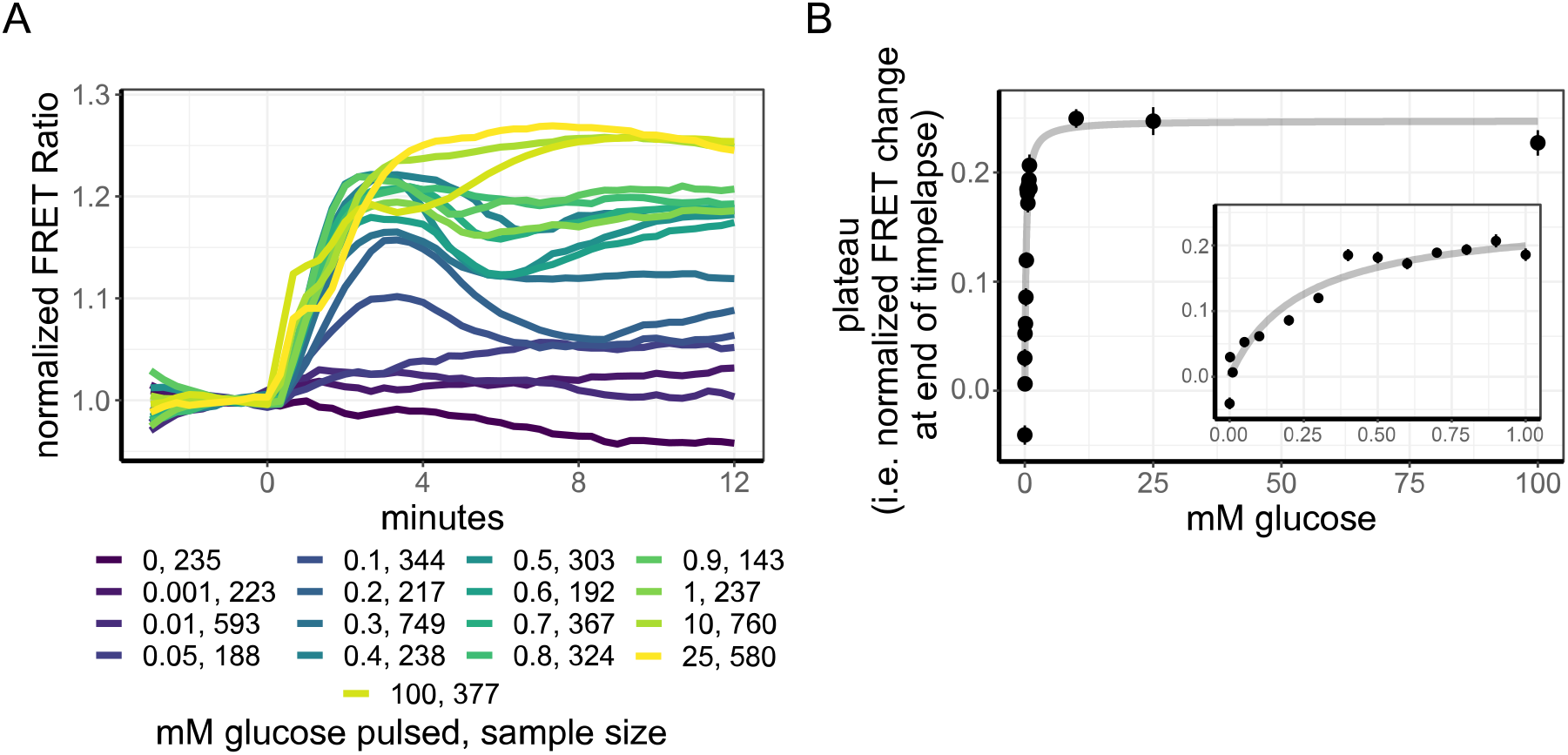
Dose-response of AKAR3-EV to glucose addition shows a high affinity. A) dose-response curves of 1% ethanol-grown W303-1A cells expressing AKAR3-EV to various concentrations of glucose additions, added at t=0 minutes. Lines show mean baseline-normalized FRET response. Colors indicate the final glucose concentration after glucose addition. B) Final FRET response of AKAR3-EV expressing cells to the glucose concentration added, dots show mean response, errorbars indicate SD, grey line indicates fit (from equation 1). Fitting of the final AKAR3-EV FRET levels to the glucose concentration pulsed shows saturation kinetics with a K_0.5_ of 0.24 and a maximal normalized FRET value of 1.24. Inset shows the zoom-in of the AKAR3-EV responses from 0 to 1 mM of glucose additions. All data obtained from at least 2 biological replicate.

In conclusion, we found that the affinity of the RRXT phosphorylation system (i.e., TORC1-Sch9 and cAMP-PKA) has a high affinity for glucose – in comparison, the affinity we found for cAMP peak height was 3.0 mM, ten times higher, therefore. Furthermore, this system appears to be homogeneous since no clear non-responders or subpopulations were found.

### AKAR3-EV shows no growth-rate dependent FRET status

After characterizing the TORC1-Sch9 and cAMP-PKA signalling response during nutrient transitions, we studied the steady state phosphorylation levels during (balanced) growth on different carbon sources. W303-1A cells expressing either the AKAR3-EV or the AKAR3-EV-NR sensor were grown to mid-exponential phase on 1% Ethanol (growth rate of 0.16 h^−1^), galactose (growth rate of 0.25 h^−1^), glucose (growth rate of 0.36 h^−1^) or mannose (growth rate of 0.34 h^−1^) and the FRET level distributions across a population was measured using a flow cytometer (Fig. 5A). We found significant differences between all conditions tested (Wilcoxon signed-rank test, P < 0.01), except between galactose and mannose (Wilcoxon signed-rank test, p-0.6). However, we also found such differences in the non-responsive sensor and no clear relation was found between growth rate and the FRET levels of AKAR3-EV after correction for aspecificity (Figs. 5B and 5C).

**Figure 5.**
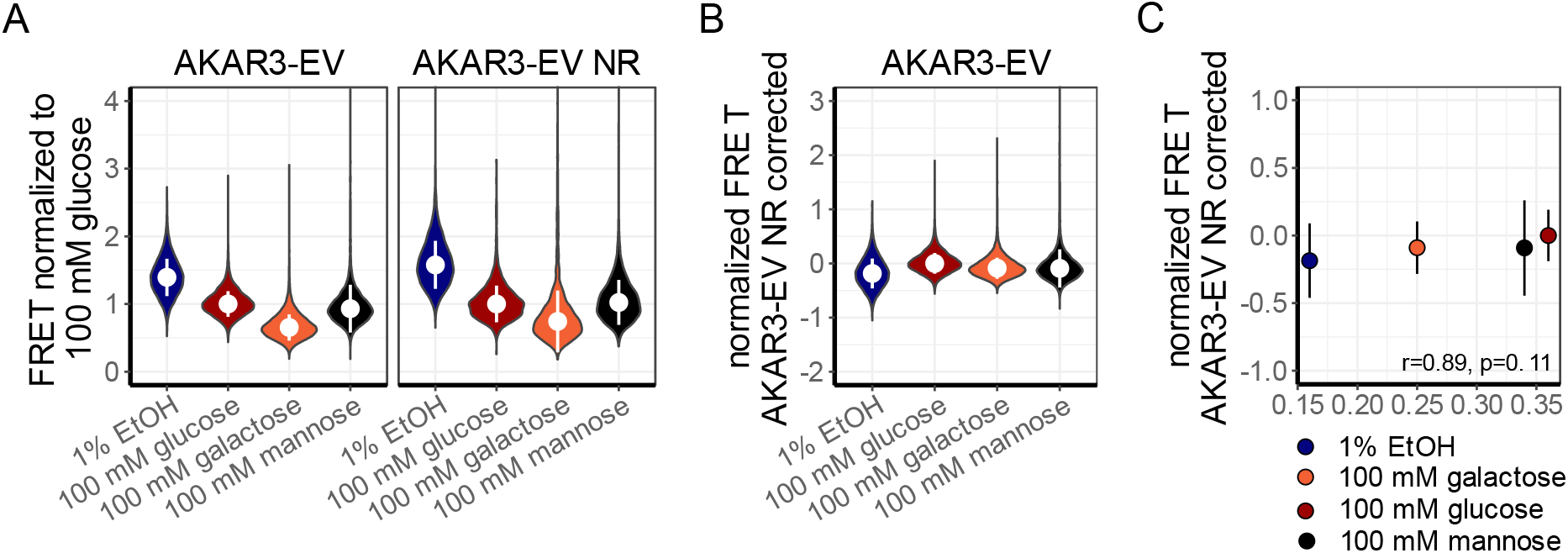
Growth rate does not relate with AKAR3EV basal levels. A) Violin plot of basal AKAR3EV and AKAR3EV NR FRET levels in W303-1A WT on various carbon sources, normalized to the 100 mM glucose condition. White dot depicts median FRET value, errorbars indicate SD. B) Violin plot of AKAR3-EV FRET levels after correcting for aspecific signal on the 4 carbon sources. White dot depicts median FRET value, errorbars indicate SD. C) AKAR3-EV FRET levels (corrected by AKAR3-EV NR signal) plotted against the growth rate of W303-1A WT on the carbon sources. Each dot depicts the population average FRET level on a specific carbon source (indicates by the dot color), errorbars indicate SD. R and p value are obtained from a Pearson’s correlation test. Representative dataset shown from 2 biological replicates.

## Discussion

In the present study we implemented the mammalian-optimized AKAR3-EV sensor in yeast and tested whether this sensor can also be applied to study Sch9 and PKA signalling in yeast. In yeast, the sensor has sufficient expression and a homogenous distribution in cells. Furthermore, growth rates were not affected by the sensor, indicating the sensor is harmless to cells (Fig. S1C). The AKAR3-EV sensor showed the expected response to an ethanol to glucose transition and outperformed a previously published AKAR sensor in yeast (Fig. 1A). We also made a non-responsive version of AKAR3-EV by mutating the RRXT motif to RRXA^59^. This AKAR3-EV NR sensor indeed showed significantly lower responses although we found that some transitions evoked a rather large response immediately after the transition. The origin of this response is not known and may be caused by nonspecific phosphorylation of the sensor domain, although this domain does not contain any other known phosphorylation sites of PKA or Sch9. This response can also originate from changes of the intracellular composition (e.g. changes in redox potential, pH, ion concentration). If needed, the AKAR3-EV NR can be used to correct for these aspecific responses. This sensor can be potentially improved for use in yeast by using phosphorylation sites that are more common for yeast (i.e., RRXS)^59^ although this could also increase the basal phosphorylation status of the sensor, decreasing its dynamic FRET range. In addition, (y)mTq2 could be used instead of eCFP since this fluorescent protein is a better FRET donor^60,61^.

The AKAR3-EV FRET response is indeed dependent on both TORC1-Sch9 and cAMP-PKA signalling (Fig. 2). Glucose addition to ethanol-grown cells showed that impaired PKA signalling (using the Tpk1^wimp^ strain) decreased the rate of phosphorylation, although the maximal response remains similar. An Sch9 deletion, on the other hand, resulted in a lower response after a glucose transition. Yet, these transitions were performed in different strains (SP1 and W303-1A, respectively). Interestingly, we also found that the W303-1A Cyr1^K1876M^ mutation (in which the transient cAMP peak is absent^13^) showed a decreased maximal response. Further research should clarify whether and why missing the transient cAMP peak has long-term effect on cell signalling status and fitness. The S25-31C strain with impaired signalling in both Sch9 as PKA, showed a small decrease in RRXT phosphorylation, proving that the AKAR3-EV sensor indeed measured solely PKA and Sch9 activity. This was further confirmed by adding rapamycin in SP1 WT and SP1 Tpk1^wimp^ cells, where the rapamycin-treated Tpk1^wimp^ strain showed a severe reduction in FRET response after glucose addition.

A major strength of biosensors is the ability to measure single-cell responses. Therefore, we assessed single-cell responses during transitions from ethanol-grown cells to glucose, sucrose, fructose and mannose (Fig. 3). As previously found for cAMP signalling^13^, hardly any heterogeneity or subpopulations were found for the glucose, sucrose and fructose transitions. In contrast, we did find a heterogenic response upon mannose addition. Mannose is known as an antagonist of cAMP signalling^13,58^ but is metabolized at rates similar to glucose as cells grow comparable on these sugars (growth rate of 0.34 h^−1^ for mannose and 0.36 h^−1^ for glucose). Glycolytic startup, determined by pH measurements also indicate that mannose is transported and metabolised at least as fast as glucose (Fig. S4). The conflicting signals between the signalling and metabolism of mannose may be the reason for the heterogenic response. We found that cells having a low basal FRET level (i.e. the FRET level before mannose addition) seem to have a lower FRET response after the shift to mannose. However, where the lower FRET levels originate from and the consequence of this heterogenic response is beyond the scope of this study and remains to be determined in future research.

The AKAR3-EV sensor revealed a relatively high affinity (K_0.5_ = 0.24 mM) of the phosphorylation system for glucose (Fig. 4). This is far below the affinity of the high-affinity hexose transporters in yeast (with a lowest K_m_ of approximately 1-2 mM, for HXT6 and 7^62,63^), which are expressed in these ethanol-grown cells. Furthermore, the affinity is lower compared to our previously obtained glucose affinity of the cAMP levels in yeast^13^, which confirms that the RRXT phosphorylation status of the cell is not solely determined by cAMP-PKA signalling. The high affinity of the RRXT phosphorylation system is, on the other site, not too far from the Monod constant for glucose-limited growth, which is around 0.5 mM, admittedly for another strain (CEN.PK)^64^. Moreover, these values are in line with previous observations for Mig1 translocation after glucose additions^65^. The fact that we did not find clear subpopulations at any glucose concentrations shows that, at least for glucose, yeast cells sense its concentration in a highly accurate and robust manner.

One large unanswered question is whether the basal signalling status of the PKA-Sch9 pathway depends on the specific growth rate. This may be expected as transcription of ribosomal genes is an important target of the pathway, and ribosomal content does scale with growth rate^66^. The obtained flow cytometry data are statistically significantly different between most conditions but given the small size of the difference relative to the spread of the distributions (Figs. 5B and 5C), their biological relevance is disputable. Therefore, we believe that the basal phosphorylation status of the RRXT motif is unchanged between various growth rates. We hypothesise that the RRXT phosphorylation response upon sugar transitions steers cells to the right cellular physiology after which this signalling system is “turned off” again. The dynamics of Fig 3A also suggest a temporal impact of sugars on the phosphorylation state.

In summary, the AKAR3-EV proved to be a robust sensor to measure the nutrient-induced RRXT phosphorylation status in yeast cells. However, since the phosphorylation status is an integrated output of the activity of both Sch9 and PKA, but also on phosphatases, its interpretation is more challenging than for other sensors. Nonetheless, we believe that the AKAR3-EV sensor is a useful addition to the toolbox that can help to elucidate how yeast cells respond and adapt to nutrient changes.

## Data and resources

AKAR3-EV and AKAR3-EV NR in pDRF1-GW can be acquired via Addgene (https://www.addgene.org/182533/ and https://www.addgene.org/182534/). Data can be found via 10.17632/w655db7rj9.1.

## Acknowledgements

We are grateful to Dr. Kazuhiro Aoki (Kyoto University) for sharing AKAR3-EV with us. We thank Dr. Sonia Colombo (Università Milano Bicocca) for providing AKAR3 in pYX212. We also thank Dr. Johan Thevelein (KU Leuven) for sharing various yeast strains.

## Supplements

**Figure S1.**
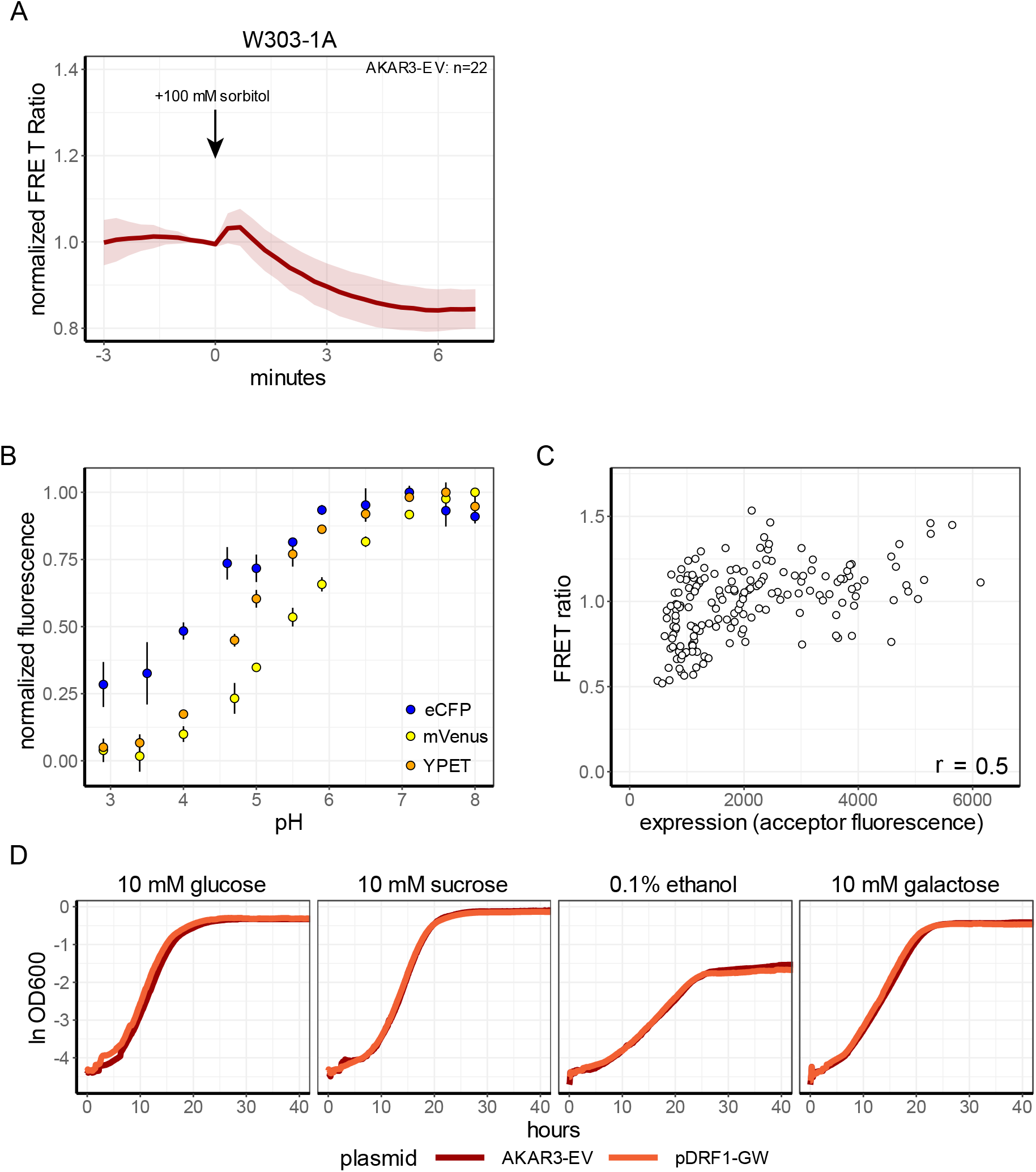
Characterization of AKAR3EV. A) AResponse of W303-1A WT cells expressing AKAR3-EV NR grown on 1% ethanol to 100 mM sorbitol addition. B) In-vivo characterization of pH sensitivity of eCFP, mVenus and YPET. Dots show mean fluorescence value, per FP (depicted by color) and pH normalized to the pH giving the highest fluorescence. Errorbars show SD. Data obtained from Botman et al., 2019^57^. C) Effect of expression levels of AKAR3-EV in W303-1A grown on 1% Ethanol on the AKAR3-EV FRET levels, each dot depicts a single cell. D) Growth curves of W303-1A WT expressing either AKAR3-EV or the empty plasmid on glucose, sucrose, ethanol or galactose. Lines show median OD value, colors indicate the expressed plasmid. All data obtained from at least 2 biological replicates.

**Figure S2.**
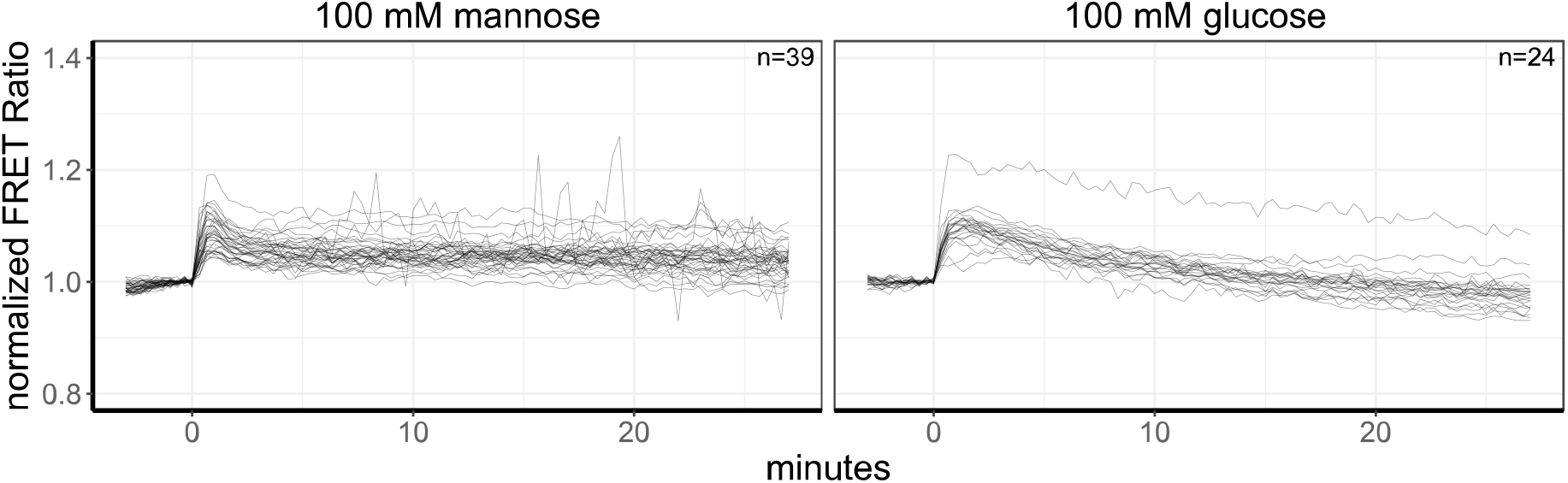
Single-cell traces of AKAR3EV-NR to mannose and glucose addition. W303-1A cells expressing AKAR3EV-NR were grown on 1% Ethanol and 100 mM mannose (left panel) or 100 mM glucose (right panel) was added at 0 minutes. Lines show single-cell traces of the baseline-normalized FRET values. All data obtained from at least 2 biological replicates.

**Figure S3.**
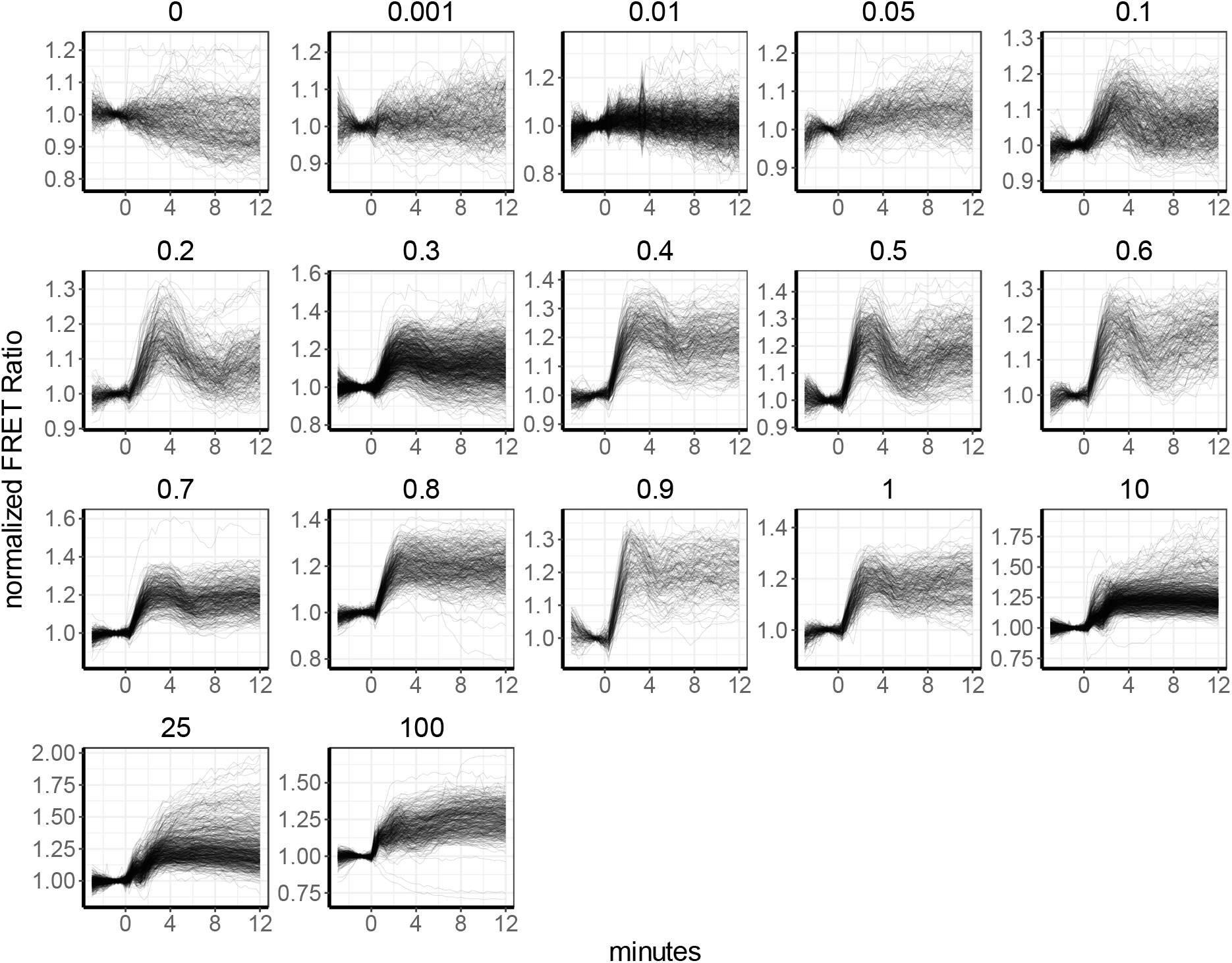
Single-cell traces of the dose-response assay. W303-1A cells expressing AKAR3EV were grown on 1% Ethanol and various glucose concentrations were added. Facet titles show the end concentration of the added glucose (in mM). Lines show single-cell traces of the baseline-normalized FRET values. Glucose was added at t=0. All data obtained from at least 2 biological replicates.

**Figure S4.**
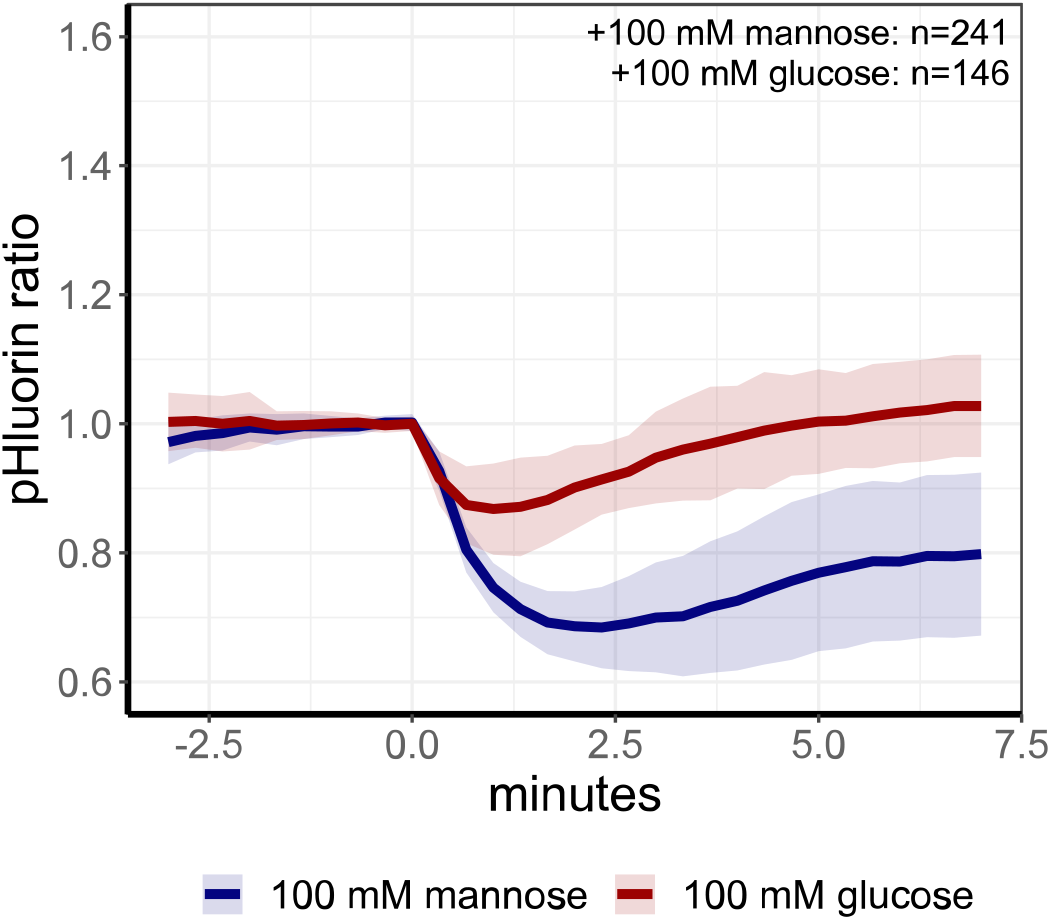
pH dynamics during glycolytic startup after glucose or mannose addition. W303-1A cells expressing pHluorin were grown on 1% Ethanol and either 100 mM glucose or mannose was added at t = 0 minutes. Lines show mean pHluorin response, colour indicates the pulsed carbon source, shades indicate SD. All data obtained from at least 2 biological replicates.

